# Impaired representation of temporal statistics in cerebellar degeneration

**DOI:** 10.64898/2026.09.12.751189

**Authors:** Tianhe Wang, Tanvi Thummala, Devika Narain, Richard B. Ivry

**Affiliations:** Department of Psychology, University of California, Berkeley, California; Department of Neuroscience, University of California, Berkeley, California; Princeton Neuroscience Institute, Princeton University, Princeton, New Jersey; Department of Molecular and Cell Biology, University of California, Berkeley, California; Donders Center for Neuroscience, Radboud University, Nijmegen, The Netherlands; Department of Neuroscience, Erasmus Medical Center, Rotterdam, The Netherlands

**Author notes:** Corresponding author : Tianhe Wang.

**Keywords:** Timing, Cerebellum, Bayesian perception, Spinocerebellar ataxia, Serial dependence

## Abstract

To increase perceptual precision, the brain is hypothesized to perform Bayesian inference, evaluating sensory input within the context of the statistical structure of the environment. The cerebellum is a strong candidate for this computation in interval timing: its integrity is essential for precise timing and supports encoding temporal statistics in animals. However, how the sensory input is combined with prior knowledge within this circuit remains unknown. To examine this, we tested patients with cerebellar degeneration and matched controls on duration reproduction tasks with varying prior distributions. Patients showed the signatures of Bayesian inference, but with greater timing variability, stronger regression toward the mean, and slower updating of the prior. We developed a model of the cerebellar microcircuit in which Purkinje cell plasticity over a granule cell temporal basis set provides a shared substrate for learning the prior and performing the inference. Simulations of cerebellar degeneration showed an output approximated a Bayesian observer with an intact prior and a noisier likelihood. This model provided a good fit of the patients’ behavior over a wide range of parameter space. These results suggest a critical role for the human cerebellum in Bayesian inference, supporting optimally timed behavior even under extensive neuropathology.

## Introduction

Accurate timing is essential for a wide range of motor and cognitive functions, such as coordinating movement and anticipating when future events will occur. However, intervals of time must be inferred from noisy sensory signals^1–3^. One prominent theoretical framework proposes that the nervous system leverages the statistical properties of the environment to improve the precision of this inference by integrating the current sensory input with prior knowledge following Bayesian principles, attenuating the effects of perceptual noise to produce an optimal estimate (Fig. 1a)^1,4,5^. The Bayesian framework has proven useful for accounting for the performance of human and non-human species across a range of cognitive and perceptual tasks^1–8^. In timing, these behavioral signatures of Bayesian inference are strikingly robust, appearing across species, sensory modalities, and interval ranges. A central question, then, is where and how the nervous system implements the requisite computations for Bayesian inference in the time domain.

**Figure 1.**
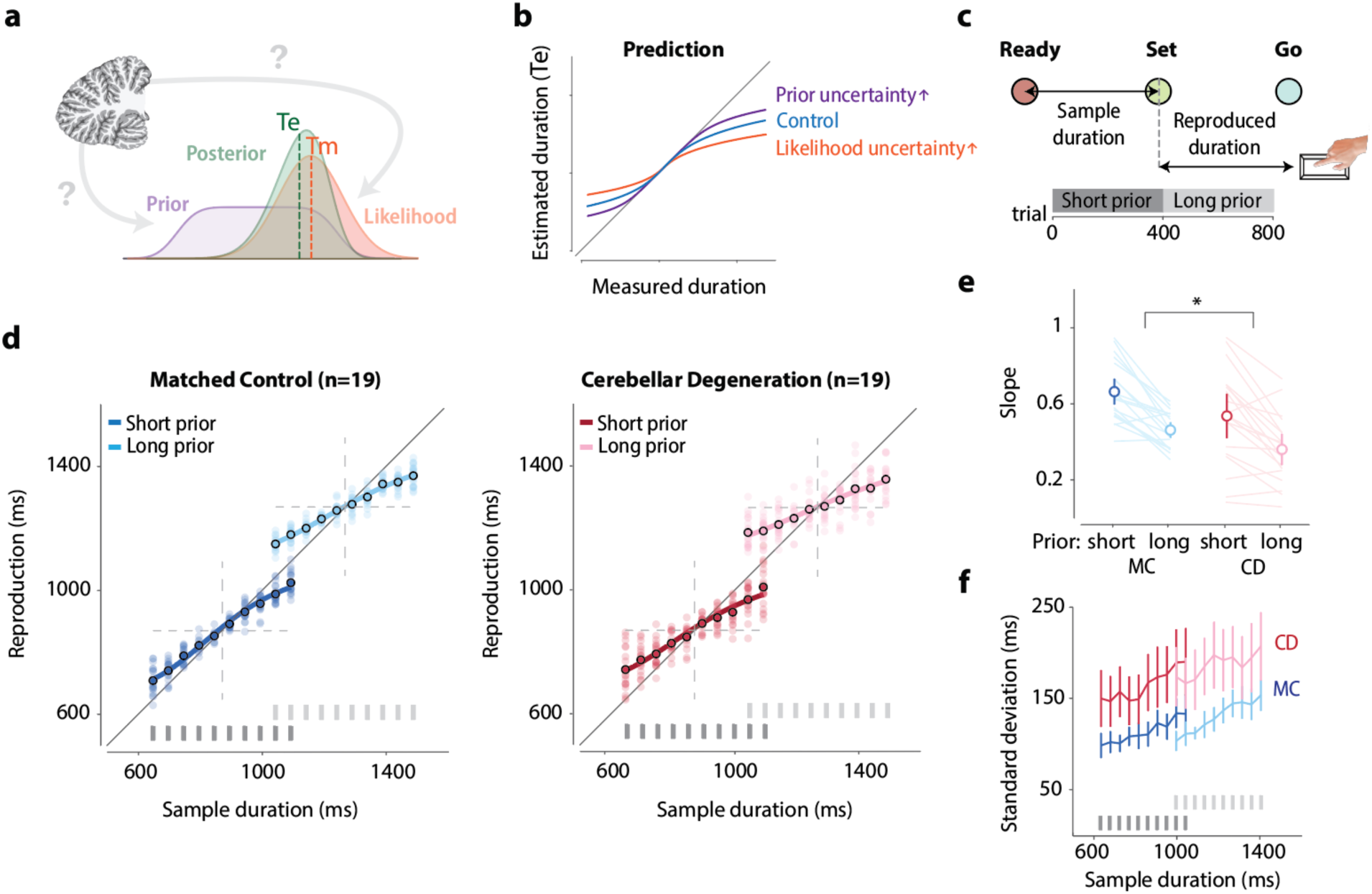
CD patients exhibit a stronger central tendency in a temporal reproduction task. (a) Schematic of the Bayesian observer model for duration perception. The nervous system forms a likelihood function of the measured duration (Tm) and integrates it with the prior distribution to yield a posterior. The estimated duration (Te) is taken as the mean of the posterior. (b) Competing hypotheses concerning how cerebellar pathology could influence central tendency: increased noise in the likelihood increases the central tendency effect, while increased noise in the prior decreases the central tendency effect. (c) Schematic of the Ready-Set-Go task. Three colored circles were presented sequentially. The sample duration was defined by the interval between the “Ready” and “Set” signals. Participants pressed the response key, attempting to synchronize their response with the onset of the “Go” signal. The interval between the “Set” signal and the key press defined the produced interval. Bottom: A short prior (range 650–1100 ms, dark gray) was employed in trials 1–400 and a long prior (range 1050–1500 ms, light gray) was employed in trials 401–800. (d) Reproduced duration as a function of sample duration for MC (left) and CD (right) across both prior conditions. Light dots represent individual trial data and black circles represent the group mean. Thick lines represent the Bayesian Least-Squares (BLS) model fit. Gray bars at the bottom indicate the sample durations that formed the short (dark) and long (light) prior distributions. (e) The slope of the function relating reproduced duration to sample duration was lower in the CD group, evidence of a stronger central tendency effect. Error bars represent 95% confidence intervals and are the same for all graphs below. *, p < 0.05. (f) Standard deviation of reproduced durations as a function of sample duration. CD exhibited greater variability compared to MC.

The cerebellum is widely implicated in behaviors that require precise timing^9–13^. Patients with cerebellar pathology exhibit increased spatial and temporal variability in their movements, a defining feature of cerebellar ataxia^14^. This variability is particularly evident on experimental tasks that directly assess the ability to produce timed movements^10,11^. Importantly, the impairment extends beyond motor production: patients with cerebellar disorders also show elevated discrimination thresholds on perceptual tasks that require precise timing, an impairment absent on non-temporal perceptual tasks^10,15,16^. More broadly, a large body of work using perturbation and physiological methods in humans and animal models has established an essential role for the cerebellum in temporal processing across diverse contexts. Eyeblink conditioning has provided a paradigmatic model given that the conditioned response (CR) is most adaptive when it is precisely timed to anticipate the unconditioned stimulus (airpuff)^17,18^. Experimental and theoretical work has highlighted the cerebellar cortex as a critical locus for precise temporal representations^19–21^. For example, lesions of the cerebellum prior to training prevent acquisition of this CR, whereas lesions of the cerebellar cortex in trained animals leave the response intact but selectively disrupt its temporal precision^12,22^.

How to characterize the role of the cerebellum in timing within the perspective of Bayesian inference remains a critical question. Central to the Bayesian framework is the distinction between two components: a likelihood function, reflecting the noisy representation of the current sensory input, and a prior distribution, capturing accumulated knowledge of the statistical structure of the environment^1,3^. These are combined to form a posterior that can shift the estimate toward the mean of the prior^1,23,24^. Recent theoretical work has proposed that plasticity within cerebellar circuits is critical for learning the prior, with empirical support coming from probabilistic eyeblink conditioning studies in mice^25^. Here it was shown that activity of cerebellar Purkinje cells is shaped by the prior distribution of stimulus time intervals and perturbing Purkinje cell activity disrupts the animals’ ability to generate behavior consistent with those statistics. While these results indicate that the cerebellum is sensitive to the temporal context, it remains unclear whether the cerebellum is also key to the inference process that entails integrating current evidence with a prior, the core computation of the Bayesian framework.

To study the role of the cerebellum in Bayesian inference, we tested patients with cerebellar pathology on a probabilistic time reproduction task, a paradigm in which behavior exhibits robust signatures of Bayesian inference across species^1,4,5^. Based on the discoveries from rodent cerebellar physiology^25^, one might expect that cerebellar pathology in humans primarily disrupts the representation of the prior. This would cause the patients’ estimates to rely more heavily on the likelihood, resulting in a weaker central tendency bias than exhibited by healthy controls (Fig. 1b). An alternative hypothesis is that cerebellar pathology might add noise to the representation of the current sensory input, yielding a less reliable likelihood, which would result in an increased bias. Note that increased variability in either component would produce a more variable posterior and, as such, predict the reduced temporal precision typically observed in cerebellar patients.

To discriminate among these hypotheses, we assessed performance in cerebellar degeneration patients and age-matched controls on a probabilistic duration-reproduction task^1,23^ that has been fundamental for evaluating the Bayesian timing theory. Consistent with prior reports, the patients were more variable in their production responses. Critically, they showed a stronger central tendency bias than controls, a result that suggests a noisier likelihood rather than a degraded prior. However, this simple interpretation is puzzling given the rodent physiology results suggesting that the cerebellum encodes prior distributions of time intervals^25^.

To resolve this puzzle, we developed a circuit-level computational model for Bayesian timing based on the architecture and physiology of the cerebellar cortex. We simulated cerebellar pathology as a reduction in the dendritic arborization of Purkinje cells, one of the most salient changes observed in patients with cerebellar degeneration^26–30^. Importantly, the model implemented a transformation that approximates Bayesian inference at the behavioral level without explicitly positing separate internal representations of the prior and likelihood. A reduction of computational capacity degraded both the likelihood and the learned prior. However, the output of the lesioned circuit behaved approximately like a Bayesian observer with an intact prior and a noisier likelihood, producing the elevated variability and stronger central tendency bias observed in patients. Taken together, we offer a mechanistic account of how the human cerebellum encodes environmental statistics to support temporal inference.

## Results

### Cerebellar degeneration increased central tendency bias

To study the influence of cerebellar pathology on Bayesian temporal inference, we employed the Ready-Set-Go (RSG) task (Fig. 1c), testing patients with cerebellar degeneration (CD, n = 19, see Table S1 for demographic information) and age-matched controls (MC, n = 19). Participants were asked to fixate on a central point while three visual cues (“Ready,” “Set,” and “Go”) were presented sequentially, each separated by the duration of the sample interval. They were instructed to measure the interval between the “Ready” and “Set” signals and then reproduce the duration by pressing a key in synchrony with the anticipated onset of the “Go” signal.

Crucially, to examine how the cerebellum contributes to the representation of prior knowledge, we manipulated the statistical context across two blocked conditions: an initial block of 400 trials in which sample durations were uniformly distributed from 650 to 1100 ms, followed by a second block in which the distribution was shifted to 1050–1500 ms (increment size 50 ms in both blocks). By comparing behavior across these two blocks, we could assess how well participants tracked the changing temporal statistics.

Previous studies of cerebellar pathology have documented increased timing variability on motor and perceptual tasks^10,11^; however, how cerebellar degeneration affects the learning of temporal statistics has remained unclear. MC exhibited the classic central tendency bias (Fig. 1d, left), overestimating shorter durations and underestimating longer ones, a hallmark of Bayesian inference. Given the physiological evidence implicating the cerebellum in learning temporal statistics, we would expect the CD group to show a weaker central tendency bias, reflecting a degraded prior.

However, the results were clearly at odds with this prediction: CD displayed a stronger central tendency effect than the MC (Fig. 1d, right). A typical way to quantify this effect is by comparing the slopes relating produced duration to sample duration across groups (Figs. 1e and S1). The slopes were shallower in the CD group relative to the MC group (2 × 2 mixed ANOVA, F(1,36) = 4.9, p = 0.033), reflecting a stronger regression toward the prior mean. Moreover, this central tendency effect was stronger for the long prior than for the short prior (F(1,36) = 59.4, p < 0.001), consistent with previous reports^1,2^. As observed here, longer durations are associated with greater variability (F(1,36) = 14.2, p < 0.001), increasing the reliance on the prior and thus strengthening the attraction toward the prior mean. The interaction between group and prior context was not significant (F(1,36) = 0.33, p = 0.57), indicating that the larger central tendency effect in the CD group was consistent across both prior distributions. Beyond the central tendency bias, CD patients also exhibited greater response variability (F(1,36) = 14.1, p < 0.001, Fig. 1f) and RMSE (F(1,36) = 13.7, p < 0.001, Fig. S2), consistent with previous studies in which patients were asked to generate a consistent duration within a block^10,11^.

To rule out the possibility that the stronger central tendency bias in CD patients reflects attentional deficits rather than a change in temporal inference, we conducted a control experiment in which we manipulated the attentional load in healthy young adults. Although reproductions were more variable under dual-task conditions, the central tendency bias was unaffected (Fig. S3), suggesting that attentional differences do not account for the CD group effect.

### The CD group showed a slower updating rate of the prior

After establishing a stronger central tendency bias in the CD group, we asked whether the rate at which the prior is updated trial-by-trial also differs between groups. Previous work has suggested that serial dependence, which describes the tendency for the reproduction of a current interval to be attracted toward the preceding one^31–33^, reflects the trial-by-trial updating of the temporal prior in the RSG task^34^. That is, if the system incorporates the current observation into the prior, the reproduction of the next interval will be biased toward the preceding sample duration (Fig. 2a–b). Based on this, we quantified the trial-by-trial attractive effect by calculating how each reproduction deviated from the mean reproduction for a given sample duration and plotted this as a function of the difference between the previous and current sample duration (Δ Sample duration).

**Figure 2.**
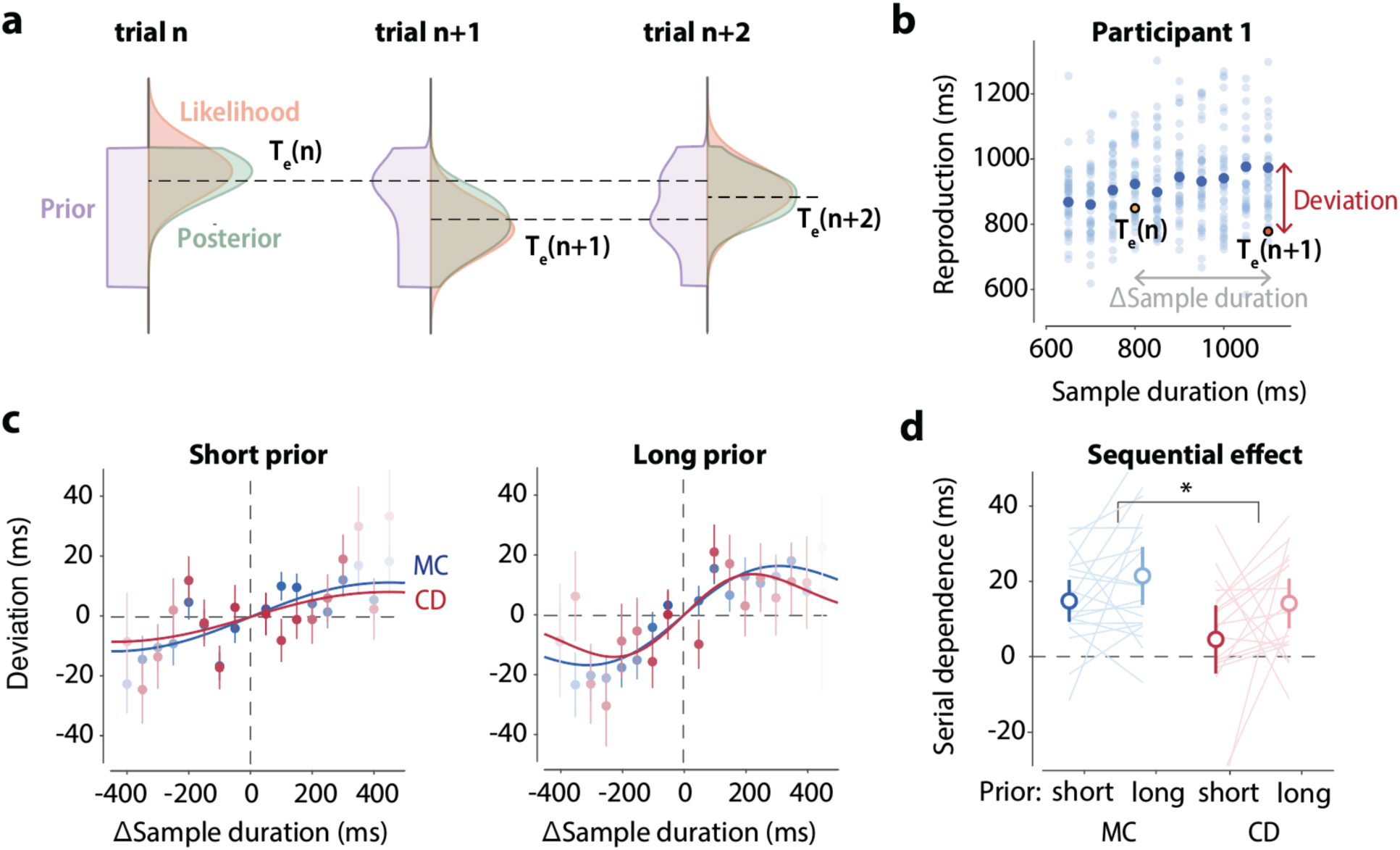
CD patients exhibit a weaker serial dependence bias. (a) Schematic illustrating how the prior is updated by T_e_ after each observation, producing serial dependence: the estimate of Te on trial *N*+1 is attracted toward the sample duration on trial *N*. (b) Reproduced duration as a function of sample duration for a representative participant. Light blue dots show responses on individual trials. Dark blue dots show the mean response for each sample duration condition. Deviation is defined as the difference between an individual response and the condition mean. Δ Sample duration is defined as the difference between the previous and current sample durations. (c) Serial dependence measured as the relationship between Deviation and Δ Sample duration for the short (left) and long (right) prior conditions in CD (red) and MC (blue). The positive correlation indicates that the reproduced duration on trial *N* is attracted toward the sample duration on trial *N*−1. Dots show mean deviation for each group, binned by Δ Sample duration. Solid lines are fits of the derivative of a Gaussian, a function commonly used to describe serial dependence. (d) Strength of the serial dependence effect, quantified as the difference in mean deviation between trials with positive and negative Δ Sample duration (see Methods). Lines show individual participants within each group; open circles show the group mean. The bracket indicates the between-group comparison. *, p < 0.05.

Both groups exhibited a positive serial dependence effect, indicating the ongoing integration of recent observations into the prior. When the previous duration was shorter than the current sample, participants tended to reproduce a shorter duration than average, and vice versa (Fig. 2c). Importantly, this attractive effect was smaller in the CD group compared to the MC group (F(1,35) = 6.1, p = 0.019, Fig. 2d), and this difference was consistent across both prior contexts (F(1,35) = 0.01, p = 0.94). The reduced serial dependence in the CD group indicates that cerebellar pathology slows the trial-by-trial updating of the prior, an independent behavioral signature of impaired prior learning.

In summary, the increased central tendency in CD would suggest that the patients have a relatively stronger impairment in their representation of the likelihood compared to the prior, a finding that is, at least superficially, at odds with current accounts of the role of the cerebellum in learning temporal statistics^25,35^. More importantly, an impairment of the likelihood alone would predict greater reliance on the prior and thus a stronger serial dependence, the opposite of what we observed. To address this puzzle, we turned to a model based on cerebellar circuitry.

### A cerebellar model for Bayesian timing

To better understand the role of the cerebellum in Bayesian inference in the time domain and how this is impacted by cerebellar degeneration, we employed a physiologically inspired cerebellar model of Bayesian timing^25,35,36^. The model focuses on the cerebellar cortex, with Purkinje cells (PCs) performing the core computations. The large dendritic arbor of the PCs receives inputs from a vast number of granule cells via parallel fibers (PFs; Fig. 3a), which convey signals originating in the brain and body to the cerebellum via the pontine nuclei^37,38^. We assume that the “Ready” cue activates a temporal basis set of granule cells, each with a Gaussian-shaped activation profile that tiles elapsed time and, as a set, exhibits scalar variance such that tuning width increases with duration (Fig. 3b). As such, the basis set corresponds to likelihood functions with increasing temporal uncertainty^39–41^. We further assume that the “Set” cue activates climbing fiber input to the PCs, an input thought to convey a teaching signal. Coincident activation of PFs and the climbing fiber induces long-term depression (LTD) at the corresponding PF–PC synapses, with the degree of LTD determined by the timing and activity level of each PF synapse relative to the climbing fiber signal (Fig. 3c).

**Figure 3.**
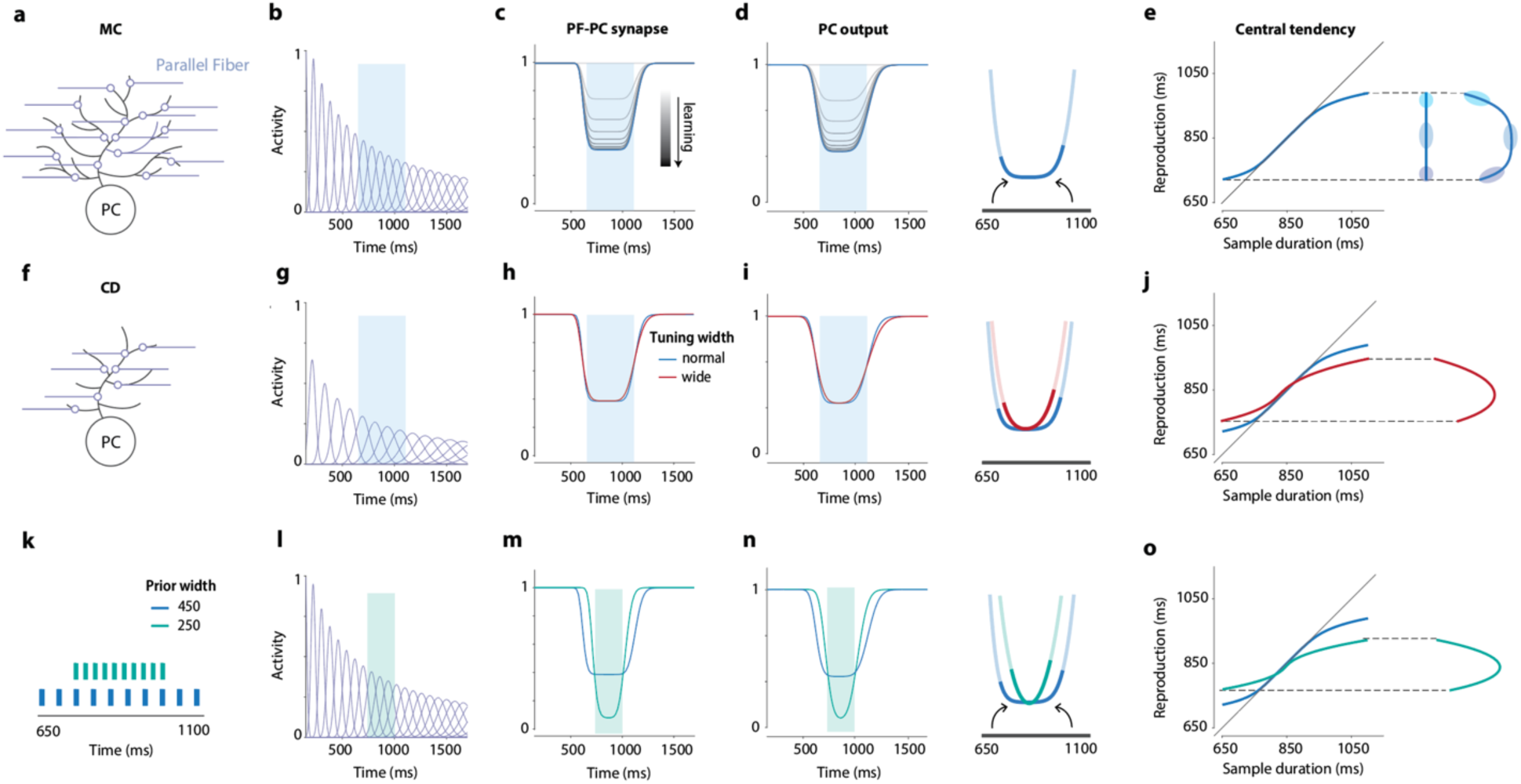
A cerebellar model for Bayesian timing. (a) Illustration of a Purkinje cell (PC) receiving input from numerous granule cells via parallel fibers (PF). (b) The temporal basis set resulting from granule cell inputs firing at different times in response to the Ready signal (0 ms). The activation profile of each granule cell follows a Gaussian distribution, with the standard deviation increasing proportionally to the peak activation time (scalar property). (c) The strength of PF–PC synapses before and after learning the prior distribution. PFs activated around the times within the prior are gradually suppressed during learning through LTD. (d) Population activity of PCs as a linear combination of granule cell inputs weighted by PF–PC synapse strength. Activity is suppressed around the range of the prior, forming a U-shaped neural trajectory (blue). The right panel zooms in on the population activity encompassing the prior. The dark blue region indicates the prior mapped onto the neural trajectory. (e) Linear readout of the U-shaped neural trajectory produces a central tendency effect in reproduced durations. Three shaded circles illustrate how compression at the two ends of the prior reduces variability, mirroring a key prediction of the Bayesian inference model. (f) Cerebellar degeneration, modeled as a reduction in the size and span of the dendritic arbor of PCs, results in fewer parallel fiber inputs and thus an impoverished temporal basis set. (g) Illustration of a 30% reduced basis set, where the remaining basis set functions exhibit greater standard deviation for each element compared to the full basis set to maintain coverage of the temporal range. (h) Broadening of the basis set elements results in a less precise representation of the prior, with information loss and exaggerated smooth transitions at the edges of the prior in the PF–PC synapse strength profile. (i) The Purkinje cell population output exhibits stronger curvature as a consequence of basis set impoverishment and increase in the width of the tuning functions. (j) The stronger curvature in Purkinje cell output produced a stronger central tendency after a linear readout. (k) Illustration of a narrow prior (750–1000 ms) compared to the standard prior (650–1100 ms). (l–o) Same as panels (g–j) but for the comparison of the narrow prior versus the standard prior.

Interestingly, this simple synaptic learning rule provides a mechanism for embedding the prior distribution in the output of the PCs^35,36^. Because intervals within the learned prior distribution are experienced more frequently, PF inputs associated with those intervals undergo stronger synaptic depression. Assuming that PC simple-spike output reflects the weighted summation of PF inputs, the resulting PC activity can be described as the convolution of this prior-dependent synaptic depression profile with the temporal basis set. This produces a U-shaped population response, in which activity is suppressed for intervals within the prior distribution and relatively preserved for intervals outside that range (Fig. 3d).

To connect this population response profile to Bayesian timing, we draw on a geometric account of Bayesian integration^36^. In duration-reproduction tasks, prior statistics have been shown to reshape low-dimensional trajectories of frontal cortical population activity^5^. Here, our model of the cerebellar cortex provides a circuit-level mechanism for generating a similar geometric configuration, where the U-shaped PC response profile defines the transformation from sampled duration to estimated duration. Specifically, suppression of PC activity within the prior range bends the representation of elapsed time through neural state space, such that equal increments in physical time no longer correspond to equal distances along the neural trajectory. When this curved trajectory is projected back onto the time axis, durations near the edges of the interval range are compressed toward the center (Fig. 3e), reducing variability in these regions and generating the central tendency bias. Thus, the model reproduces two signatures of the Bayesian Least-Squares (BLS) estimate^1^. Moreover, although the two edges of the prior distribution are symmetric in probability, they produce an asymmetric depression profile in PC output due to the broadening of the temporal basis functions for longer durations. This asymmetry predicts stronger central tendency effects for longer, more uncertain intervals, consistent with behavioral observations in humans and other species^1,4^.

The curvature of PC output is governed by two factors, the encoding precision of the granule cell basis set (i.e., the width of the likelihood functions) and the shape of the prior distribution itself. Increasing the width of the granule cell tuning functions (broader likelihoods) would enforce a low-pass filtering while embedding the prior into the synaptic weights of the Purkinje cell arbor (Fig. 3g–h). This would lead to attenuated encoding of peripheral intervals represented in the prior and, therefore, would cause added compression in the PC population output at the two ends of the prior. When the readout is linear, increased compression in this population representation will result in a stronger central tendency effect (Fig. 3i–j). As such, when switching from short to long prior, the model will predict a stronger central tendency as now the tuning width of the basis set functions increases due to the scalar property (Fig. S4).

The width of the prior also shapes the curvature of PC output. When the prior becomes narrower, the same prior probability is concentrated over a smaller range of intervals, producing stronger and more localized LTD at the corresponding PF–PC synapses. This sharper prior-dependent depression accentuates the curvature of PC output at the boundaries of the prior, so that the linear readout again produces stronger regression toward the prior mean (Fig. 3k–o).

In sum, this model offers a mechanistic link between single-neuron plasticity in the cerebellar cortex and a population-level representation that implements Bayesian-like integration. All of the model predictions regarding the changes in likelihood and prior range are consistent with normative Bayesian theory and empirical results observed in studies of temporal processing with healthy participants^1,2,24^.

### Modeling the effect of cerebellar degeneration on Bayesian timing

How can we model the effect of cerebellar pathology within this framework? As shown in humans and mouse models, the most prominent feature of cerebellar degeneration is the loss of Purkinje cells^42,43^. This typically begins with dendritic atrophy^26–30^, reducing the number of PF–PC connections. Within the context of our model, a consequence of this pathology would be an impoverishment of the basis set input to the Purkinje cells. Based on work demonstrating that cerebellar granule cell representations remap to maintain coverage of the relevant temporal range^39^, we assume that the tuning functions of the spared granule cells expand in width, preserving coverage of the full interval range. This would come at a cost in temporal precision (Fig. 3f–g). As noted above, reduced precision in the basis set functions produces a stronger central tendency effect (Fig. 3h–j), consistent with the finding that the CD patients showed an enhanced central tendency effect (Fig. 4a). To further illustrate this principle, we ran a series of simulations with varying degrees of dendritic loss to mimic the effects of a variable disease process; we assumed that the width of the tuning functions increases as an inverse function of the lesion size. The slope of the central tendency effect decreased monotonically as dendritic loss increased for both prior conditions (Fig. 4c).

**Figure 4.**
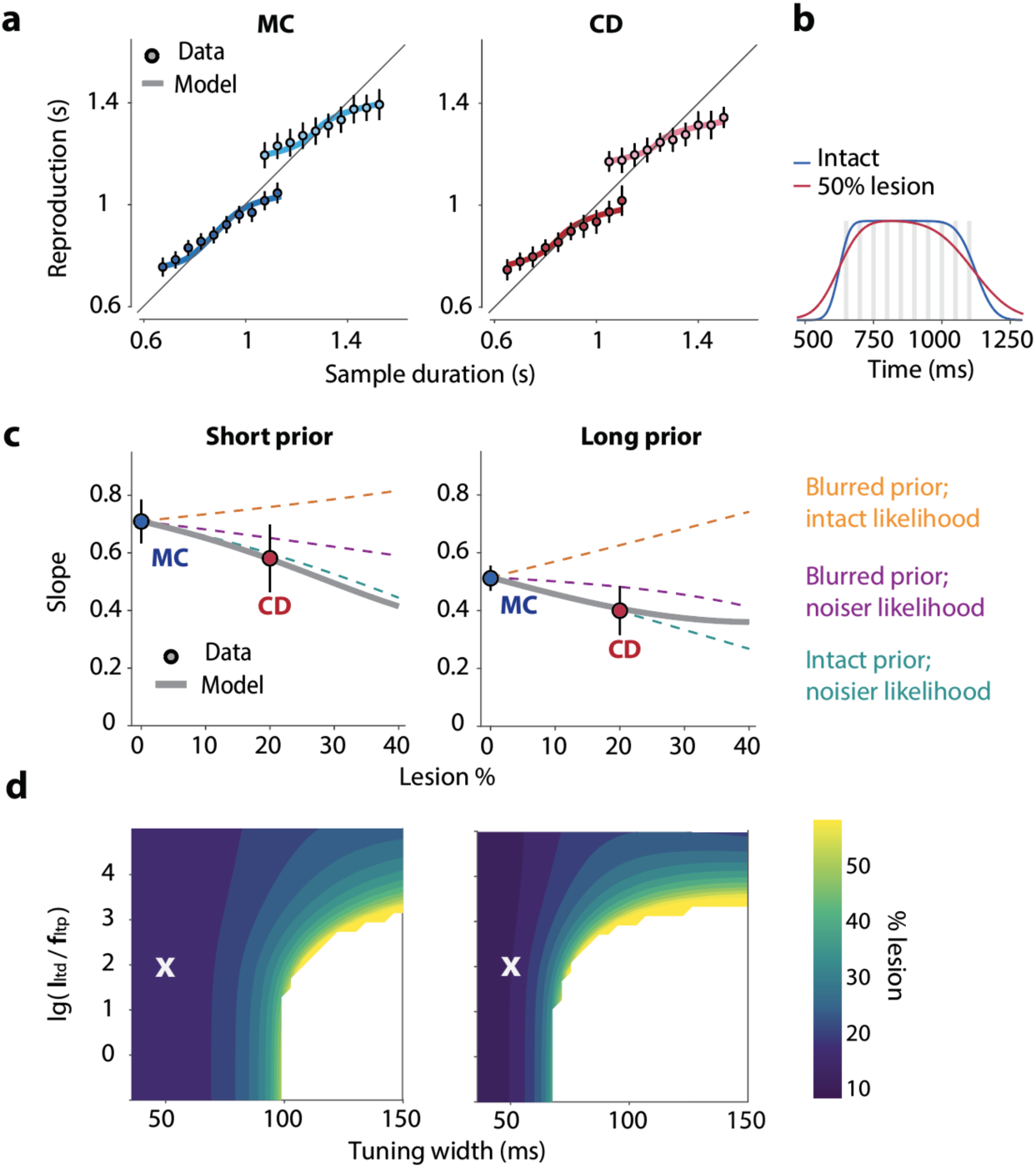
Reduced precision of the granule cell basis set quantitatively accounts for the enhanced central tendency effect in cerebellar degeneration. (a) Prediction of the best-fit model alongside empirical data for the reproduced duration. MC are shown in blue and CD in red with dark and light shades denoting the short and long priors, respectively. (b) Blurred measurement of the short prior. Convolution of the prior (gray bars indicating ten sampled durations) with the granule cell tuning functions, shown for the intact basis set (blue) and after a 20% lesion (red). (c) Slope of the central tendency effect as a function of lesion size for both prior ranges, predicted by the circuit model (gray) and by three normative Bayesian observers: 1) An intact likelihood with a prior blurred by the lesion (orange); 2) A noisier likelihood with the blurred internal prior (purple); 3) A noisier likelihood with an intact prior (teal). For each BLS observer and prior range, the width of the likelihood before the lesion was fitted so that the observer’s slope matched the intact model. (d) Minimum lesion size required to reproduce the observed change in central tendency effect for different combinations of basis set standard deviation and ratio of the learning and forgetting rates. White regions indicate parameter combinations with no solution. The cross indicates the parameter combination used in panels (a–c). The x axis depicts the standard deviation of an element within the basis set that has a mean of 1000 ms. *l_ltd_* and *f_ltp_* represent the learning rate and forgetting rate, respectively.

We next examined whether the model could quantitatively predict the performance of the MC and CD groups. We fit the model to the group-averaged reproduction functions of both groups simultaneously. The best-fit model indicated that a loss of approximately 20% of dendritic connections could account for the increase in the central tendency effect observed in CD (Fig. 4c). Moreover, the model accurately predicted the reproduction functions of both groups across the two priors (Fig. 4a). To assess the robustness of the model predictions, we varied two additional parameters, the baseline width of the tuning functions in the basis set and the LTD/LTP ratio (Fig. 4d). The model successfully predicted the observed change in central tendency across a wide range of values for both parameters, indicating that the predicted link between dendritic loss and amplified central tendency reflects a robust model property rather than a fit to specific parameter choices. The model failed when the tuning function width was very large and the LTD/LTP ratio was low. However, in this parameter regime, the suppression of PF–PC synapses was too weak to capture the prior shape even in the non-lesioned model (healthy participants), and is therefore unlikely to be biologically relevant (Fig. S5).

It is important to note that unlike the canonical Bayesian model, in which the prior and likelihood are independent, the precision of the prior representation in the cerebellar model is directly constrained by the resolution of the basis set (Fig. 4b), the neural population that encodes the sensory measurement. Increasing the width of the basis set functions will broaden the likelihood function and simultaneously degrade the representation of the prior. Critically, inference in this circuit does not require the integration of separate representations of the prior and the likelihood because the two are already combined in the learned synaptic weights. The blurred prior (Fig 4b) is simply the distribution of the circuit’s own measurements, and the estimate can be read out directly from the Purkinje cell output. Indeed, across lesion sizes, the model mimicked a Bayesian observer with an intact prior and a noisier likelihood (Fig 4c, teal).

We recognize that there are alternative ways to model cerebellar pathology within this general framework. Dendritic atrophy could result in a sparse basis set in which tuning functions are unevenly distributed due to missing connections, with the width of the residual tuning functions unchanged (Fig. 5a). Gaps in the basis set within the prior range create dips in the PC output function (Fig. 5b). The distortion in the shape of the PC output function resembles what is observed when the prior is narrowed (Fig. 3n). Critically, this model of temporal scotomas also results in a stronger central tendency effect compared to an intact model (Fig. 5c–d).

**Figure 5.**
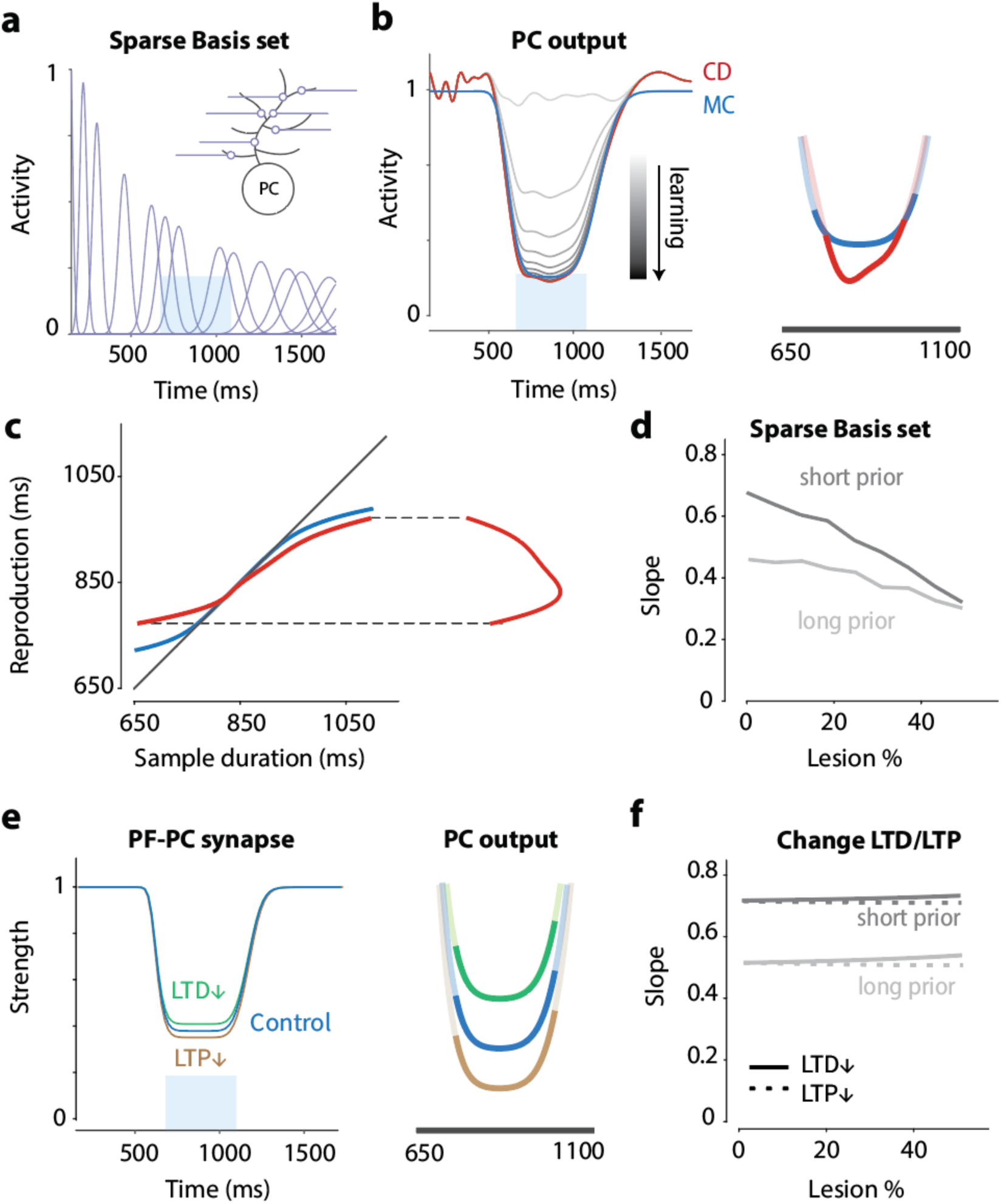
Alternative models of cerebellar degeneration. (a) Illustration of an alternative way to model cerebellar degeneration. As in our initial model, degeneration is assumed to reduce the dendritic arbor of Purkinje cells. However, in this alternative, no remapping of the basis set is applied, leaving uneven coverage of the temporal space. (b) Left: The sparse basis set results in inhomogeneous baseline activity (light gray) when averaged across a population of Purkinje cells, and an imperfect representation of the prior after learning (red). Right: Zooming in on the population activity over the prior range shows that missing tuning units distort the shape of the output function, resembling what is observed when there is greater suppression of the basis set functions. (c) Readout from the simulated PC trajectory in panel b, showing the enhanced central tendency effect in reproduced duration. (d) Predicted slope of the central tendency as a function of the proportion of elements lesioned from the basis set, for the short (dark) and long (light) prior. (e) Left: Changes in learning dynamics (ratio of LTD to LTP) influence the degree of suppression of parallel fiber–Purkinje cell (PF–PC) synapses. Right: While the absolute level of PC activity differs across conditions, changes in the LTD/LTP ratio have minimal effect on the curvature of the final neural trajectory. (f) Predicted slope of the central tendency as a function of lesion size under reduced LTD (solid) or reduced LTP (dotted), for the short (dark) and long (light) prior. Changes in the LTD/LTP ratio have little influence on the slope.

Moreover, as suggested by the serial dependence results, cerebellar degeneration may also influence learning dynamics. One factor that influences Purkinje cell output during the learning of the prior is the ratio between LTD (learning) and LTP (forgetting) rates. In principle, decreasing the LTD/LTP ratio results in a shallower representation of the prior and increased curvature of the Purkinje cell output, thus producing a stronger central tendency. Conversely, increasing the LTD/LTP ratio would weaken the central tendency effect. However, we found that varying the LTD/LTP ratio over a biologically relevant range (0.5– 2 fold) had minimal influence on the slope and, as such, the impairment of LTD or LTP could not account for the observed magnitude of change in the empirical data (Fig. 5e–f).

Together, these results indicate that decrease the encoding power within the granule cell basis set, rather than altered learning dynamics, provides a parsimonious account of the stronger central tendency effect observed in the CD group.

## Discussion

A central question in neuroscience is how contextual information is acquired and represented by the brain^6,44^. Bayesian inference has provided a powerful framework for understanding how context shapes performance across a wide range of perceptual and cognitive tasks. One such domain is time perception, where behavioral evidence of Bayesian inference has been observed across species^1,5,45^. However, insights into the underlying neural mechanisms remain scarce, particularly in humans. To address this, we combined behavioral testing in patients with cerebellar degeneration with a circuit-level model rooted in cerebellar anatomy and physiology. Specifically, we examined how cerebellar pathology affects the acquisition and use of temporal priors in a probabilistic duration reproduction task. Both patients and matched controls showed the hallmarks of Bayesian timing, including the central tendency effect and serial dependence, indicating that their estimates were shaped by a temporal prior^1,24,46^. The CD group, however, exhibited greater overall variability in their reproduced intervals, a stronger central tendency bias, and weaker serial dependence. To account for these results, we developed a model of the cerebellar cortex. Importantly, the likelihood and prior are not separable in this model. Rather, the activation pattern across the basis set elements constitutes the likelihood^47,48^, with the prior embedded in the synaptic weights between these elements and the Purkinje cells. Readout of the Purkinje cell population produces the basic features of Bayesian timing. When the basis set is impoverished, a simulation representing cerebellar degeneration^26–30^, temporal representation and the prior are both degraded, yet the output of the circuit approximates a Bayesian observer operating with an intact prior and noisier likelihood, showing increased response variability and a stronger central tendency effect. These results provide an account, at the level of a defined neural circuit, of how the human brain represents the temporal statistics of the environment and exploites that information in inference.

Our study brings together several lines of research relevant to the role of the cerebellum in timing. Human neuropsychological studies established that cerebellar damage inflates the variability of perceiving or reproducing a single duration without shifting the mean estimate^10,15^. With the emergence of Bayesian models^1,24,46^, psychophysical studies in both humans and rodents showed that the temporal context modulates perceived duration through the formation of a prior. Many frameworks have been proposed for how the brain encodes prior knowledge, and temporal context in particular^5,35,36,49–51^, with theoretical studies suggesting a critical role for the cerebellum in learning the prior^25,35^. However, the supportive evidence has been limited to studies with mice using paradigms that require only that the animal learn the distribution of the stimulus set in order to time their response. A Bayesian observer, by contrast, must not merely represent the temporal context but apply it in an inference process, estimating the interval by integrating the current likelihood with the prior to form a posterior.

To bridge this gap, we provide, to our knowledge, the first assessment of this process in humans, testing a population in which cerebellar function is compromised. Interestingly, the CD group was more variable in reproducing intervals and showed a stronger central tendency bias. Under a classic Bayesian model, this pattern would be taken to indicate that cerebellar damage degraded the measurement of elapsed time (the likelihood) and, thus, greater weight was given to an intact prior^25^. On this account, the cerebellum is seen as supplying the likelihood, with the representation of the prior involving extracerebellar structures. However, further evidence from the serial dependence results argues against this hypothesis. Because the attraction toward the preceding interval reflects the influence of a recently updated prior, a noisier likelihood should strengthen serial dependence along with the central tendency. Instead, we observed that serial dependence was weaker in the CD group.

An important insight arising from our model is that one need not posit separate representations of the prior and the likelihood, but rather that both arise from the dynamics of activity in the cerebellar cortex^9,48,52^. Activity across a subset of the basis set elements, the granule cells, provides the likelihood, a noisy representation of the current input, while the prior is emergent from the synaptic weighting of this input onto the Purkinje cells. Lesioning the basis set degrades both representations, yielding a noisier likelihood and a less precise prior. However, what governs behavior is the readout of the Purkinje cell population. Given the geometry of the Purkinje cell activity, the lesioned circuit would still approximate a canonical Bayesian observer with a noisier likelihood and veridical prior. As such, an adaptive property of this architecture is that even in a state of neuropathology, estimation remains near-optimal, albeit operating on noisier measurements. We note that the readout mechanism, based on the the geometry of Purkinje cell activity, determines the slope of the central tendency effect; this mechanism does not capture the trial-by-trial dynamics that give rise to serial dependence. The weaker serial dependence in the CD group may reflect an additional deficit in PC plasticity, beyond the loss of PF–PC connection. Interesingly, changes in the learning and/or forgetting rate has minimal effect on the centeral tendecy effect.

Notably, the core predictions of the model are not dependent on how cerebellar degeneration is simulated. Given evidence suggesting that the cerebellum exhibits some degree of flexibility to adjust the temporal basis set in a task-dependent manner^39^, we reasoned that it would show similar flexibility to maintain coverage of the relevant temporal range as granule cells are lost. To achieve this, we assumed that the tuning functions of the spared granule cells would broaden at a cost in temporal precision. This broadening changes the shape of the Purkinje cell output function, increasing the curvature at the edges of the prior range. A consequence of this increase in curvature is that, with a linear readout of this function, there will be greater compression toward the mean of the sample distribution. An alternative would be to model degeneration as gaps in the basis set, or what might be called temporal scotomas. Under this variant, the PC output function is similarly distorted. Moreover, the circuit mechanism we describe should generalize beyond the specific form of the basis set. Recent theoretical work has shown that arbitrary basis sets can yield comparable Purkinje-cell readouts after weighted integration^53–55^, indicating that the precise form of individual granule-cell tuning is not critical. More important, the central claim, that reducing basis set precision increases the curvature of the Purkinje-cell read-out trajectory and thus amplifies temporal compression, is a geometric property that should hold for any basis set capable of supporting Bayesian-like timing.

A limitation of the current study is the heterogeneity of our cerebellar cohort, which included a mixture of genetic subtypes known to differ in the regional distribution of Purkinje cell loss and in the extent of extracerebellar involvement^42,43^. Nevertheless, Purkinje cells are among the most consistently affected populations across subtypes, reflecting the selective vulnerability of these neurons^26,27,56^. Moreover, although the affected regions vary, most patients showed elevated timing variability, indicating that the circuitry supporting temporal processing was compromised in these individuals. As such, the effects we report are unlikely to reflect the idiosyncrasies of a particular pathology, but rather represent a general consequence of cerebellar degeneration.

Whether the principles we describe here generalize beyond cerebellar timing remains an open question. As noted above, the classic Bayesian framework treats the prior and the likelihood as independent representations. In contrast, in our model the two share a substrate and are combined without an explicit integration step. Perception may work the same way. The prior for orientation perception appears to reflect natural scene statistics^57,58^ and cued expectations evoke feature-specific templates in V1, even in the absence of any stimulus^59,60^. This suggests that the prior is expressed in the same region that encodes the sensory evidence, the likelihood. Our cerebellar model offers a potential mechanism for how such statistics come to be embedded in an encoding population to support Bayesian integration. In contrast, in working memory and decision-making tasks, the two appear separable. For example, silencing the posterior parietal cortex reduces the central tendency bias without interfering with the processing of the current stimulus^61^. Determining how the brain implements these schemes, and adjudicating between them across domains, is an important direction for future work. More broadly, the current study illustrates how the statistical structure of the environment can be embedded directly within the substrate that encodes the incoming sensory evidence.

## Methods

### Participants

Nineteen individuals with cerebellar degeneration and nineteen age- and education-matched control participants were recruited for this online study. CD patients were drawn from a patient database maintained by our laboratory, which includes individuals recruited from ataxia support groups across the United States and a recruitment flyer posted on the National Ataxia Foundation website^62^. MC were recruited via advertisements circulated on internet forums. The protocol was approved by the Institutional Review Board at the University of California, Berkeley. Participants were financially compensated ($40) for their participation.

Initial inclusion in the CD group required self-report of ataxia, with final inclusion based on a detailed medical history and clinical exam consistent with cerebellar ataxia. Seventeen of the participants had genetic confirmation of their disorder (16 with an SCA subtype, one with POLR3A-related ataxia) and two patients exhibited cerebellar ataxia of unknown origin. To assess motor status, we administered the Scale for the Assessment and Rating of Ataxia (SARA)^63^ to both CD and MC. To establish general neuropsychological status, we administered the Montreal Cognitive Assessment (MoCA)^64^. Both scales were modified for online administration^62^. Demographic, diagnostic, neurological, and neuropsychological data for each individual in the CD group, along with summary information for the MC group, are provided in Table S1. The CD group showed a mean SARA score of 10.1, representing a mild-to-moderate motor impairment. There was no significant difference between the CD and MC groups on the MoCA test.

### Ready-Set-Go task

The experiment was conducted online, with participants accessing the program from their homes using personal computers. An experimenter monitored the session via a Zoom video call to provide instructions and ensure task compliance. The experimental code was written in JavaScript and administered via the Google Chrome web browser. Visual stimuli were presented on the participant’s monitor, and responses were made by pressing the spacebar on the computer keyboard. We verified that the refresh rate of each participant’s monitor was at least 60 Hz. Prior to the experiment, we performed a slow-motion video analysis on both PC and MacBook hardware to confirm that the temporal error in stimulus presentation did not exceed one refresh frame (≤ 16.7 ms) on either operating system. All data were collected and stored using Google Firebase.

We used the Ready-Set-Go temporal reproduction task^1,34^. This task is ideal for the planned Bayesian analysis since it involves the presentation of a range of target durations and the durations defined by the produced responses are continuous rather than categorical (as is the case with most perception tasks). Each trial consisted of a sequential presentation of a red, green, and blue circle (200-pixel radius) at the center of the screen. The duration of each circle was 50 ms and the critical manipulation was the stimulus-onset asynchrony. This was identical for the two intervals and varied from trial to trial (see next paragraph). The interval between the red and green circles defined the sample duration and the participant’s task was to press the spacebar simultaneously with the blue stimulus^2,65,66^. The instructions emphasized that the participant should attempt to synchronize their keypress with the onset of the blue circle rather than respond in reaction to the appearance of the blue circle. The reproduced duration was measured as the time from the onset of the second stimulus until the key press. No feedback was provided although participants could estimate their performance based on their sense of whether the blue circle appeared to precede or follow their response.

The experiment consisted of two blocks of 400 trials. In the first block, the sample duration for each trial was selected from a uniform distribution that ranged from 650 to 1100 ms. The actual sample values were separated by 50 ms (e.g., 650, 700, 750, etc.), with each of the 10 durations presented 40 times in a random order. For the second block of trials, the distribution of sample values ranged from 1050 to 1500 ms, with the actual values again separated by 50 ms. We did not provide a break between the two blocks, nor did we inform the participants of the distributional manipulation. The total time on task was approximately 75 minutes. Although no formal breaks were scheduled, participants were allowed to take self-initiated breaks as needed. All MC completed both blocks (800 trials). However, due to fatigue, we terminated the second block for some of the patients (see Table S1 for the trial number). The minimum number of completed trials was 40 for the second block, a sufficient number to estimate the strength of central tendency in the new prior.

### Behavioral analysis

For each sample duration, we calculated the mean and standard deviation of the produced intervals on an individual basis. Produced intervals shorter than 300 ms or more than 2.5 SD from the participant’s mean were treated as outliers and were excluded from further analysis (4.2% for CD and 2.7% for MC). To measure the central tendency effect, we performed a linear regression on the data from each block to calculate the regression coefficient between the reproduced durations and the sample durations. As we are mainly interested in the central tendency effect rather than systematic shifts, we realigned the averaged reproduced duration of each individual to the prior mean (875 ms and 1275 ms for the short and long prior, respectively) for illustration (Fig. 1d) and model fitting (see below).

To measure the trial-by-trial learning of the prior, we examined serial dependence effects, the influence of the previous observation on the current percept. We defined a “deviation” index as the reproduced duration for a given trial minus the mean reproduced duration of all trials with that target duration^34,67^. The deviation is plotted as a function of the change of sample duration (Δ Sample duration, trial N−1 – trial N). A positive correlation between the deviation scores and Δ Sample duration indicates that the current perception is attracted toward the previous sample duration, and is taken as a signature that the prior has been updated after the previous trial. We fit this function with a derivative of a Gaussian, a model typically employed to describe serial dependence effects^33^. To quantify the serial dependence effect, we calculated the difference between the average deviation when Δ Sample duration is positive and the average deviation when Δ Sample duration is negative^34,67^ (Fig. 2d). One CD participant whose serial dependence index was more than 3 SD above the group mean in both prior conditions was excluded from this analysis.

Group differences were evaluated with 2 × 2 mixed-design ANOVAs, with Group (CD, MC) as a between-participant factor and Prior (short, long) as a within-participant factor. For two measures, reproduction variability and RMSE, Levene’s test indicated unequal variances between the groups, reflecting the greater heterogeneity of the CD group. For these measures we confirmed the group effect with Welch’s t test and with a Wilcoxon rank-sum test on the participant means averaged across the two priors. The statistical outcomes of these two tests were concordant with the ANOVA results (variability: t(24.3) = 3.76, p = 0.001, rank-sum p = 0.003; RMSE: t(31.5) = 3.71, p < 0.001, rank-sum p = 0.005). In all tests, the significance level was set to two-tailed p < 0.05.

### Bayesian Least-Squares (BLS) Model

We applied the Bayesian Least-Squares (BLS) model developed by Jazayeri and Shadlen^1^. We assumed that the internal prior (*π*(*t_s_*)) reflects the true prior distribution used in each condition: a uniform distribution ranging from 650 to 1100 ms for the short prior and from 1050 to 1500 ms for the long prior. Following Bayes’ rule, the likelihood function models the relationship between the perceived duration (*t_p_*) and the sample duration (*t_s_*) as:

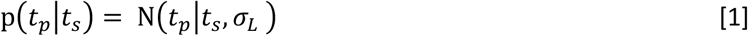

where *σ_L_* defines the perceptual noise. The posterior, π(*t_s_*|*t_p_*), is the normalized product of the prior and the likelihood function:

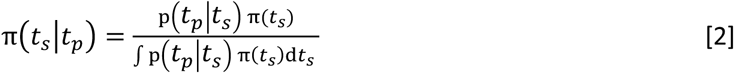

A loss function, *l*(*t_e_*, *t_s_*), was used to obtain the Bayesian Least-Squares (BLS) estimate, *t_e_*, the mean of the posterior under the assumption that it minimizes the mean squared error:

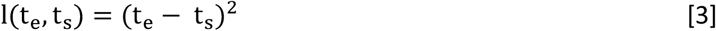

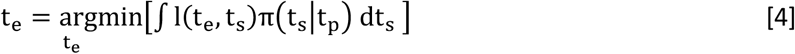

Finally, a motor response (t_r_) is made based on t_e_ and motor noise *σ_M_*.

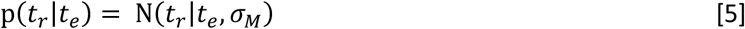

We assume *σ_M_* , *σ_L_* exhibit scalar properties: *σ_M_* = *k_M_*t_e_ ; *σ_L_* = *k_L_t_s_* , with *k_M_* and *k_L_* as the free parameters of the model. We fit the model to the group-averaged reproduction function for each prior condition by maximizing the log-likelihood using the fminsearch function in MATLAB. The posterior was estimated by numerical integration.

### A Cerebellar model for Bayesian timing

To examine how cerebellar pathology may contribute to the behavioral deficits observed in the CD patients, we employed a cerebellar timing model inspired by neurophysiological observations^35^. This model assumes that the firing rate of each granule cell *i* follows a Gaussian distribution:

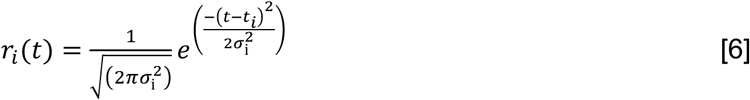

where *t_i_* is defined as the time with peak activation in granule cell *i*. *σ_i_* is the standard deviation of a Gaussian-shaped basis function which follows the scalar property: 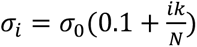, where N is the total number of granule cells and *k* is a scaling factor. The peak activation times are spaced evenly across the simulated range of elapsed times. The width of the elements in the basis set increases with the peak time of the elements. Since the area under the curve for each element is constant, peak amplitude falls with peak time. As such, elements activated later in the interval are both broader and lower in amplitude. This idealized basis set is based on normative considerations^9,25,35,53^ and is consistent with what has been observed in a recent physiological study^68^.

For the encoding of *p*(t_s_), we define *w_i_*, the strength of the synaptic connection between granule cell *i* and a Purkinje cell. For each PF–PC synapse, long-term depression (LTD) is modeled as proportional to the firing rate of the GC shortly before the firing of climbing fibers (CFs) at the time when the Set signal is presented (end of sample interval). PF–PC synapses are eligible for LTD during the eligibility trace (*ε*), a short time window for plasticity before CF firing. For simplicity, we set this time window to 0 ms relative to the Set signal onset assuming *ε* equals the transmission time for the Set signal to reach the cerebellum.

In the presence of GC firing in the absence of CF stimulation, a weak restoring force, long-term potentiation (LTP), acts to reverse the effects of LTD. The dynamics of LTD and LTP are governed by their respective time constants, *τ_ltd_* and *τ_ltp_*. As such, we have

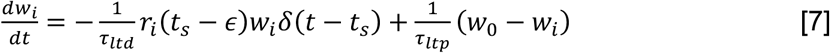

where *δ* is the Dirac delta function. This can also be expressed in a trial-by-trial manner by assuming the trial intervals are roughly the same:

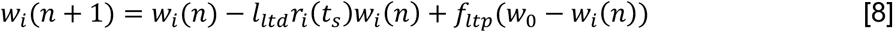

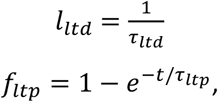

where *t* is the trial duration. As such, the synaptic strength in the stable state, *w_i_*(∞), is determined by the ratio between *l_ltd_* and *f_ltp_*.

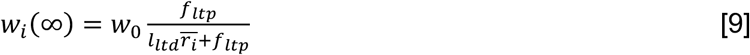

The overbar in Eq. 9 denotes the mean activation of granule cell i across the sample durations of the prior and we set w_0_ = 1. Because depression scales with the current synaptic weight, w_i_ remains bounded within (0, w_0_]. PC output was modeled as a linear combination of granule cell inputs weighted by the PF–PC synapses:

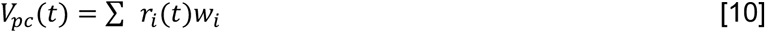

A linear readout was applied to estimate the duration *t*_e_ from *V_pc_*. To implement this, we first introduced a vertical scaling parameter, *S*, applied to the y-axis of *V_pc_* . We projected the scaled *V_pc_* onto an encoding axis (the x-axis). The readout window was defined as t ∈[*t_s_*_,*min*_, *t_s_*_,*max*_]. We aligned *V_pc_* and the readout window at the midline and then calculated the trajectory length along the *V_pc_* curve on both sides; the distance traveled along *V_pc_* was mapped to the sample duration, while the original x-axis represented the readout (perceived/reproduced duration).

### Modeling cerebellar degeneration

To simulate the effects of cerebellar degeneration, we consider two variants of basis set degradation, corresponding to two distinct assumptions about how the elements of the circuit are impacted by dendritic loss.

In the first variant, the tuning of the elements in the basis set is remapped to compensate for missing inputs. This is motivated by recent work showing that cerebellar granule cell representations can flexibly remap to maintain coverage of behaviorally relevant temporal intervals^39^. Under this view, the loss of granule cell inputs is compensated by a rescaling of the remaining basis functions such that full coverage of the interval range is preserved at the cost of reduced temporal precision. To implement this, we manipulated the number of elements in the basis set (N), reducing it by a factor corresponding to the lesion size (e.g., a 50% lesion reduces N by half). To preserve full coverage of the temporal range, we increased the standard deviation of the tuning functions (*σ*_O_), and simultaneously adjusted the vertical scaling parameter (S). By co-varying these two parameters, the baseline PC output (*V_pc_*) of the lesioned model matched that of the intact model before learning the prior distribution. Both σ₀ and S were scaled by the inverse of the factor by which N was reduced; for example, in the 50% lesion case, both were doubled. The remaining tuning functions are redistributed evenly across the temporal range, broadened in width but reduced in number, yielding the impoverished-but-uniform basis set described in Fig. 3f–g.

In the second variant, the basis set undergoes no remapping. Instead, we randomly removed a specified percentage of the basis set elements, keeping the initial width and spacing of the surviving elements. As such, coverage of the temporal range becomes uneven, with gaps where the elements are missing. This yields a sparse basis set described in Fig. 5a.

### Parameter settings and robustness tests

To simulate participant behavior, we set the size of the basis set to N = 200 elements, with mean activation times spanning 0–4000 ms. The number of elements does not influence model predictions provided it is sufficiently large to extend beyond the sampled temporal range. The free parameters (*l_ltd_*/*f_ltp_*, *σ*_O_, *k*, and *S*) were determined by a grid search over σ_0_, k and the ratio l_ltd_/f_ltp_ that minimized the difference between the predicted slopes and the group-averaged slopes. For each combination, S was calibrated so that the intact model matched the slope of the healthy participants in the short-prior condition, ensuring a reasonable baseline before any lesion was applied. The search returned a best-fitting parameter set of σ_0_ = 70 ms, k = 2.5, and l_ltd_/f_ltp_ = 100 (l_ltd_ = 10, f_ltp_ = 0.1), with a calibrated S = 7.4 × 10^5^. Models of cerebellar degeneration (Figs. 4 and 5), implemented by broadening the basis set tuning functions, reducing the number of basis set functions (Fig. 5a–d), or decreasing *l_ltd_*/*f_ltp_* (Fig. 5e–f), were evaluated by applying the lesion to this best-fit baseline model. We further performed a robustness test on our main cerebellar model (the remap variant) by systematically varying *l_ltd_*/*f_ltp_* and *σ*_O_ (Fig. 4d). For each combination, *k* and *S* were re-fitted to the healthy participants’ data so that the baseline model produced a reasonable pre-lesion prediction, and patient data were then simulated by applying the lesion to this baseline model.

### Comparison with Bayesian observers

We compared the circuit with observers using BLS estimates (Eqs. 1–4) in which the measurement noise was proportional to the width of the basis set element tuned to the sample duration, σ_m_(t_s_) = cσ₀(0.1 + kt_s_/T)/(1 − L), where T = 4000 ms, L is the lesion size and c is a constant. For the first observer (Fig. 4c, teal), it always used a veridical uniform prior with the effect of degeneration increasing the noise of the likelihood. In the second observer (orange), the likelihood variance was unaffected by the lesion and the prior was blurred by a Gaussian whose variance equaled the lesion-induced increase in measurement variance. In the third observer (purple), the noise of the likelihood increased and the prior was blurred in the same way as the second observer. For each observer and prior range, c was fitted so that the observer matched the slope of the intact model prior to the lesion to provide a fair point of comparison. The blurred internal prior in Fig. 4b was computed with c = 1, the width of the basis set itself.

## Author contributions

TW, DN, and RBI contributed to the conceptual development of this project. TT collected the data and TW analyzed the data and prepared the figures. TT and TW wrote the initial draft of the paper. TW, DN, and RBI were involved in the editing process.

## Acknowledgments

RBI is funded by the NIH (grants R35NS116883 and R01DC017091). DN is funded by the Netherlands Organization for Scientific Research (Vidi-193.076, Aspasia-015.016.012, Gravitation-024.005.022), AiNed foundation and the NGF (NGF.1609.241.021).

## Supplementary Results

**Table S1.** Demographic and neuropsychological summary of participants.

| **CD** | **Age** | **Gender** | **Type** | **SARA** | **MoCA** | **Education** | **Trials** | **MC** | **Age** | **Gender** | **MoCA** | **Education** |
|---|---|---|---|---|---|---|---|---|---|---|---|---|
| **1** | 52 | M | SCA2 | 13.5 | 25 | 14 | 800 | **1** | 49 | M | 27 | 16 |
| **2** | 57 | F | SCA28 | 5.5 | 26 | 14 | 800 | **2** | 59 | M | 27 | 18 |
| **3** | 32 | M | SCA3 | 6.5 | 28 | 16 | 800 | **3** | 53 | F | 27 | 22 |
| **4** | 55 | F | Unknown | 6 | 25 | 18 | 800 | **4** | 61 | F | 26 | 16 |
| **5** | 73 | F | SCA3 | 7 | 27 | 19 | 800 | **5** | 48 | F | 28 | 16 |
| **6** | 49 | F | SCA14 | 10 | 30 | 18 | 600 | **6** | 67 | M | -- | -- |
| **7** | 60 | F | SCA6 | 10 | 28 | 18 | 580 | **7** | 62 | F | 27 | 16 |
| **8** | 69 | F | SCA2 | 8.5 | 28 | 19 | 760 | **8** | 49 | M | 29 | 16 |
| **9** | 57 | F | SCA6 | 8.5 | 29 | 18 | 680 | **9** | 73 | F | 29 | 18 |
| **10** | 48 | F | SCA3 | 25.5 | 22 | 18 | 480 | **10** | 68 | M | 28 | 18 |
| **11** | 48 | F | SCA6 | 12.5 | 27 | 17 | 800 | **11** | 76 | F | 29 | 18 |
| **12** | 62 | F | SCA6 | -- | -- | 14 | 800 | **12** | 42 | F | 29 | 18 |
| **13** | 51 | F | Unknown | 13.5 | 26 | 18 | 800 | **13** | 77 | F | 26 | 20 |
| **14** | 51 | F | SCA6 | 14 | 29 | 16 | 440 | **14** | 47 | F | 24 | 18 |
| **15** | 72 | M | SCA6 | 10 | 27 | 18 | 800 | **15** | 54 | M | 26 | 16 |
| **16** | 38 | M | SCA6 | 7 | 26 | 16 | 800 | **16** | 48 | M | 27 | 16 |
| **17** | 59 | F | POLR3A | 10 | 30 | 21 | 800 | **17** | 74 | M | 23 | 18 |
| **18** | 39 | M | SCA28 | 8 | 26 | 20 | 800 | **18** | 44 | F | 28 | 13.5 |
| **19** | 63 | F | SCA6 | 6 | 27 | 14 | 660 | **19** | 51 | F | 29 | 18 |
| **Mean** | 54.5 |  |  | 10.1 | 27 | 17.2 |  | **Mean** | 58 |  | 27.1 | 17.3 |
| **SD** | 11.1 |  |  | 4.7 | 2 | 2.1 |  | **SD** | 11.5 |  | 1.7 | 1.9 |

**Figure S1.**
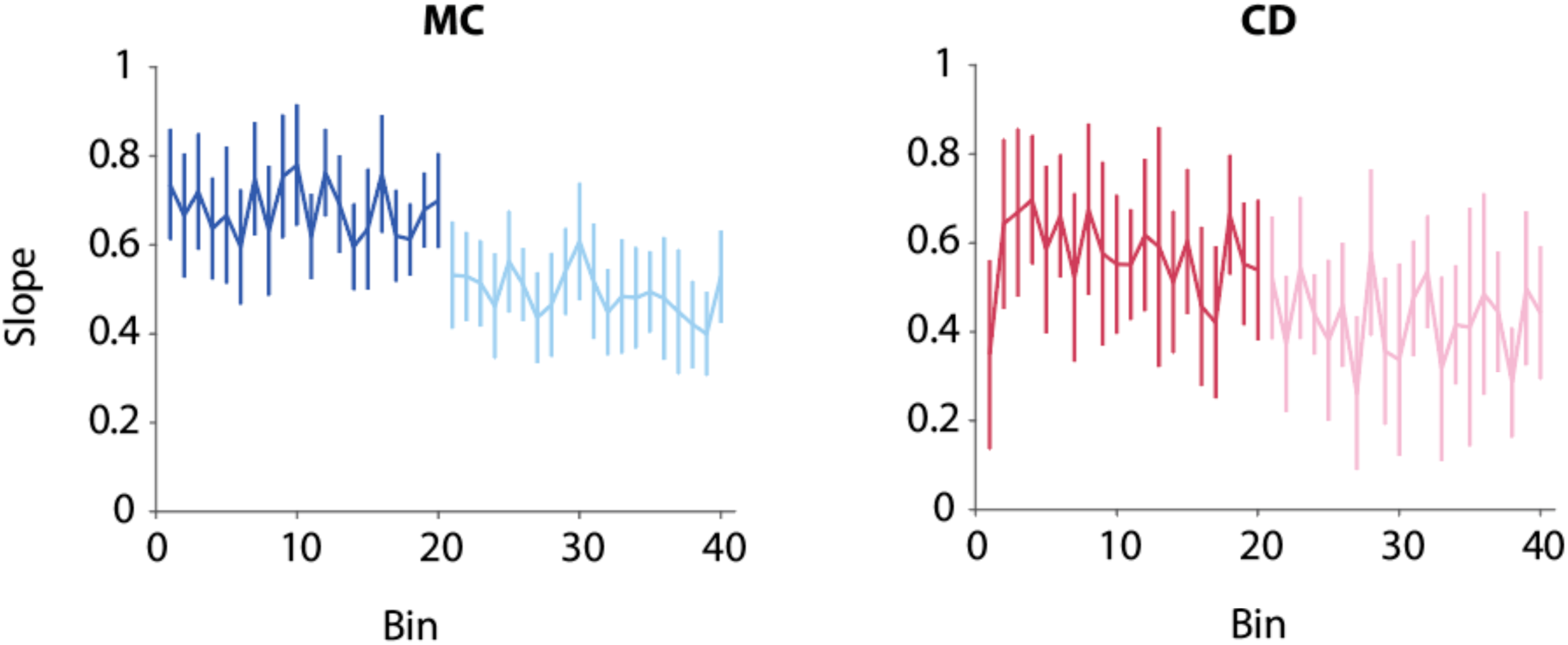
Time course of the central tendency effect. Each bin represents a block of twenty trials with each sample duration within the prior presented twice. Although the slope appears to decrease with increased experience with each prior, suggesting that the central tendency effect increases, the estimates are very noisy, especially for the CD data (red).

**Figure S2.**
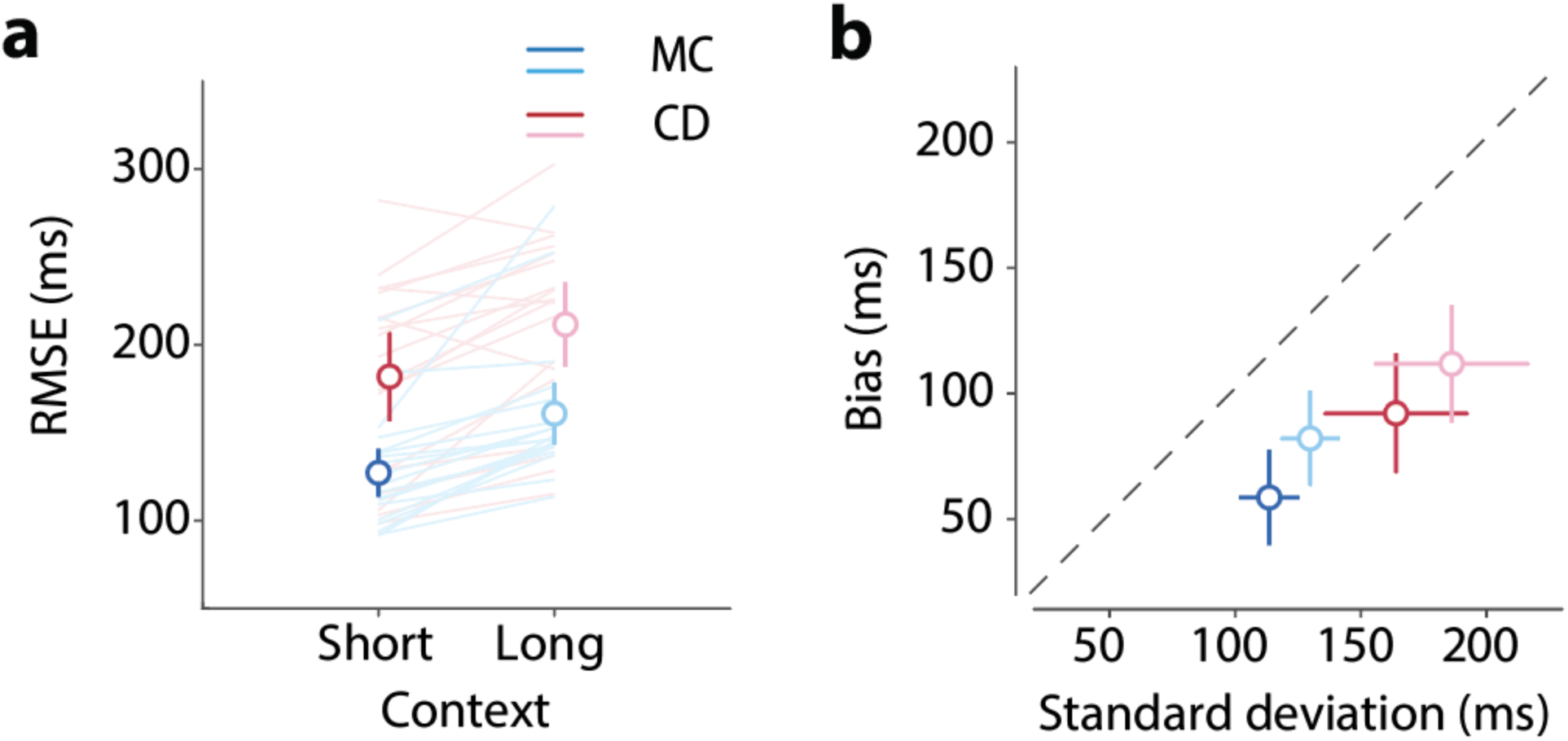
RMSE and variance of reproduced durations. (a) CD showed higher root mean square error (RMSE) compared to MC, indicating greater overall deviation from the sample durations. (b) RMSE decomposed into bias (y-axis) versus response variability (standard deviation; x-axis). The CD group exhibited both larger variability and bias (also see Fig. 1).

**Figure S3.**
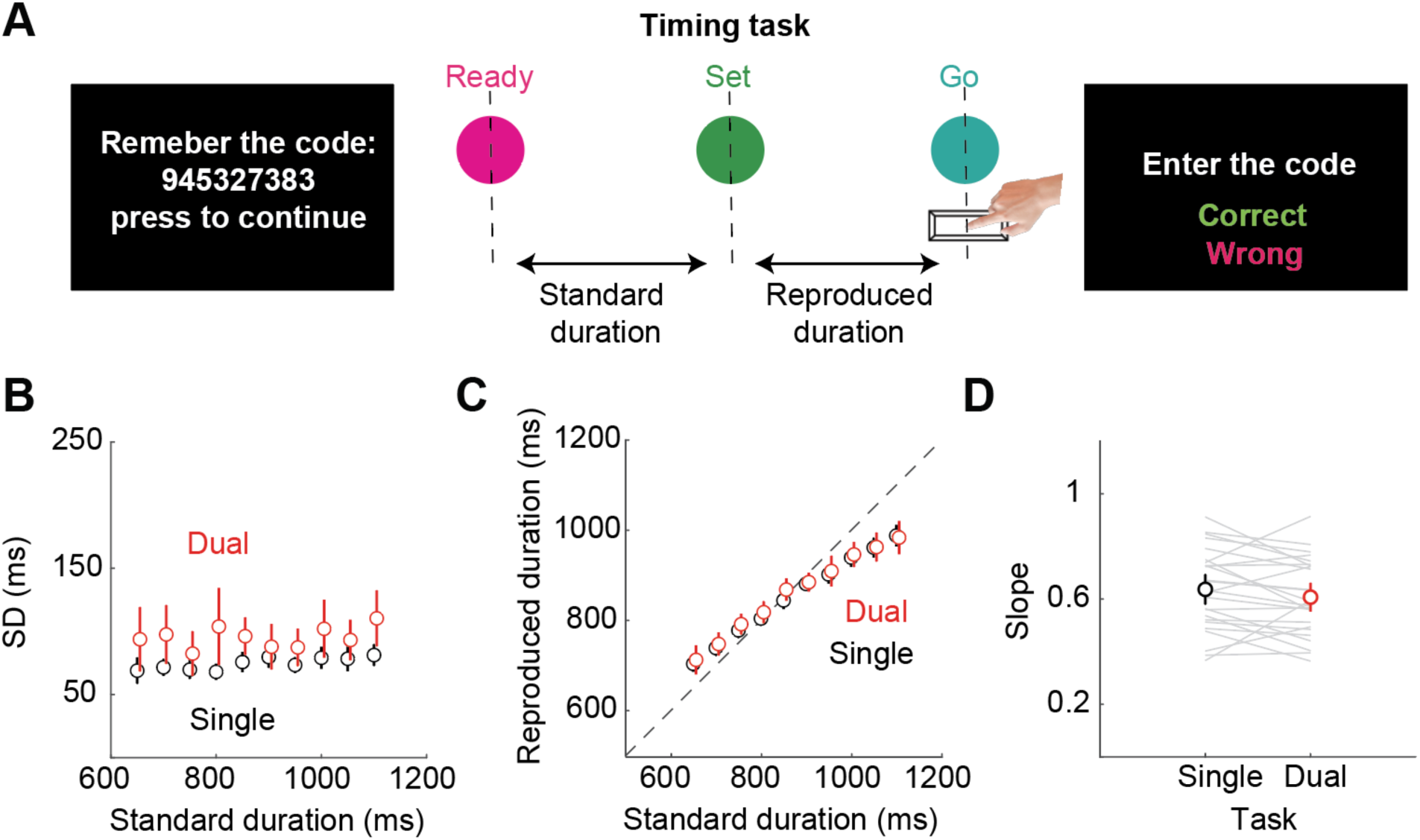
Central tendency effect is not influenced by attentional load. An alternative account of the larger central tendency effect in the CD group is that the patients fail to attend to the task on a subset of the trials and, on these trials, resort to using the prior mean. To test this hypothesis, we tested 25 healthy young adults in a supplemental experiment in which a dual-task manipulation was used to examine the effects of attention on the central tendency effect. (a) On 25% of trials, a sequence of 9 or 10 digits was presented at the start of the trial, and the participants were instructed to remember the list. When ready, the participant initiated the Ready-Set-Go task by pressing a key. One second after they pressed the response key to mark the reproduced duration, they were prompted to enter the digit sequence on the keyboard. Binary feedback (Correct/Wrong) indicated if they were correct on their report of the digit sequence. For the other 75% of trials, no digits were presented, and the participant only performed the timing task. The mean accuracy to recall the number string in the working memory task was 71.2 ± 17.8% (mean ± SD), indicating that this was a challenging secondary task. (b) The standard deviation of the reproduced duration increased under the dual-task condition, further demonstrating that the two tasks shared some common attentional resources (t(24)=4.3, p<0.001). (c, d) However, the central tendency effect was unaffected by the attentional manipulation. The average reproduced duration was nearly identical between the single- and dual-task conditions, and the slope of these functions did not differ between the single- and dual-task conditions (t(24)=1.4, p=0.16).

**Figure S4.**
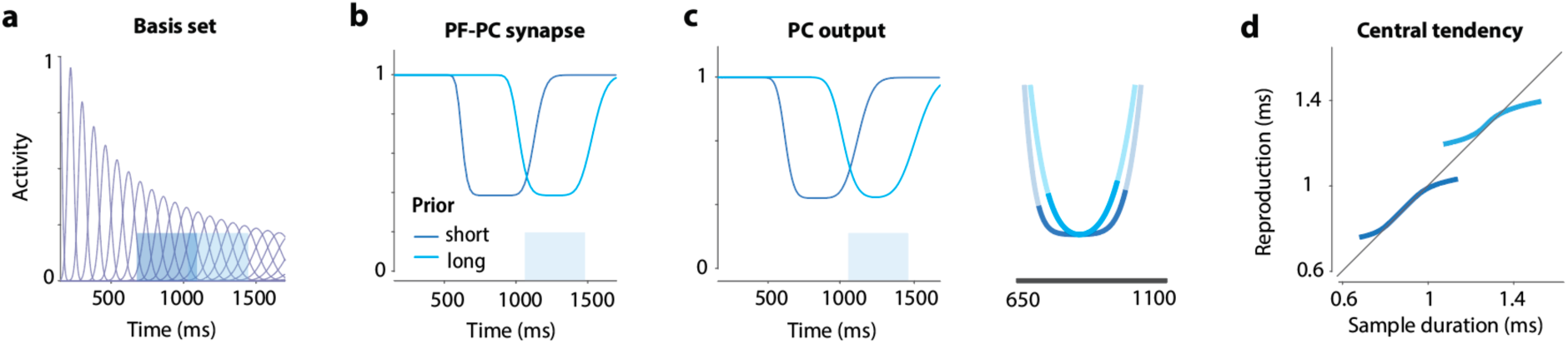
The cerebellar model predicts a stronger central tendency when the mean of the prior increases. (a) The temporal basis set. Due to scalar properties, the units tuned to the long prior (1050–1500 ms) have wider tuning functions compared to those tuned to the short prior (650–1100 ms). (b–c) The strength of parallel fiber–Purkinje cell synapses and the Purkinje cell (PC) output after learning the two priors. The wider tuning functions in the long prior result in a less precise prior representation and stronger curvature in the PC output. (d) Linear readout of the PC output shows a stronger central tendency for the long prior compared to the short prior.

**Figure S5.**
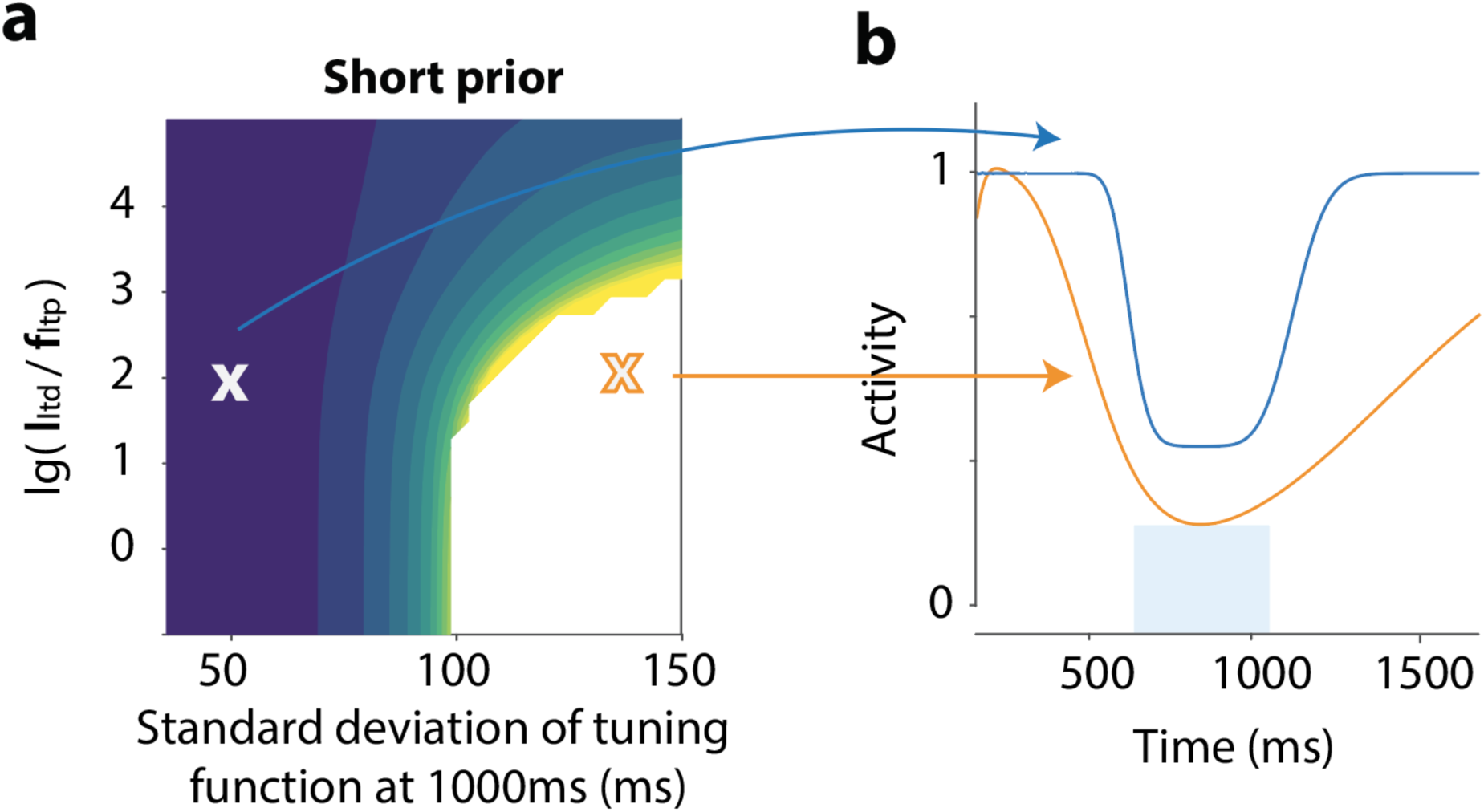
The cerebellar model fails to learn the prior when the basis set tuning function is too wide. Left panel replicates Figure 4d, showing the minimum lesion size required to reproduce the observed change in central tendency effect for different combinations of basis set standard deviation and the learning-to-forgetting rate ratio. The right panel shows the Purkinje cell (PC) output for two sample parameter sets. The blue line shows the best-fit parameters, while the orange line has the same LTD/LTP ratio but triple the basis set function width. With these parameters, the PC output function poorly reflects the shape of the prior even before lesion.

